# Protein Profiles: Biases and Protocols

**DOI:** 10.1101/2020.06.13.148718

**Authors:** Gregor Urban, Mirko Torrisi, Christophe N. Magnan, Gianluca Pollastri, Pierre Baldi

**Author notes:** Authors contributed equally to this work (Gregor Urban, Mirko Torrisi, Christophe N. Magnan). Email addresses* (Gregor Urban), (Mirko Torrisi), (Christophe N. Magnan), (Gianluca Pollastri), (Pierre Baldi).

## Abstract

The use of evolutionary profiles to predict protein secondary structure, as well as other protein structural features, has been standard practice since the 1990s. Using profiles in the input of such predictors, in place or in addition to the sequence itself, leads to significantly more accurate predictors. While profiles can enhance structural signals, their role remains somewhat surprising as proteins do not use profiles when folding in vivo. Furthermore, the same sequence-based redundancy reduction protocols initially derived to train and evaluate sequence-based predictors, have been applied to train and evaluate profile-based predictors. This can lead to unfair comparisons since profile may facilitate the bleeding of information between training and test sets. Here we use the extensively studied problem of secondary structure prediction to better evaluate the role of profiles and show that: (1) high levels of profile similarity between training and test proteins are observed when using standard sequence-based redundancy protocols; (2) the gain in accuracy for profile-based predictors, over sequence-based predictors, strongly relies on these high levels of profile similarity between training and test proteins; and (3) the overall accuracy of a profile-based predictor on a given protein dataset provides a *biased* measure when trying to estimate the actual accuracy of the predictor, or when comparing it to other predictors. We show, however, that this bias can be avoided by implementing a new protocol (EVALpro) which evaluates the accuracy of profile-based predictors as a function of the profile similarity between training and test proteins. Such a protocol not only allows for a fair comparison of the predictors on equally hard or easy examples, but also completely removes the need for selecting arbitrary similarity cutoffs when selecting test proteins. The EVALpro program is available for download from the SCRATCH suite (http://scratch.proteomics.ics.uci.edu).

## 1. Introduction

Protein structure prediction is usually decomposed into simpler but still difficult tasks like the prediction of secondary structure, relative solvent accessibility, domains, or contact/distance maps. Despite the variety of methods proposed to tackle each of these tasks, the use of evolutionary information, notably sequence profiles, in the input of the prediction methods has been a constant since the 90s when it was shown to significantly improve prediction accuracies (Rost and Sander, 1993). It is not uncommon to report an improvement of 10% or more when using profiles instead of sequences alone.

A key reason for why profiles improve accuracy is that they can amplify structural signals against the noisy background of evolution. For instance, an alternating pattern of buried and exposed residues in a profile, typically signals the presence of an alpha helix on the surface of a protein (Benner and Gerloff, 1991). The same pattern can be less visible at the level of an individual sequence. However, this observation alone does not provide a full explanation for their usefulness for two main reasons. First, proteins do not use profile information when folding in vivo. Thus, in principle, one may expect sequence-based predictors to be able to achieve the same level of accuracy as profile-based predictors; however, this has not been observed in the past 30 years. Second, over the same time period, the same sequence-based redundancy reduction protocols–initially derived to train and evaluate sequence-based predictors–have been applied to train and evaluate profile-based predictors. However, if we visualize a profile as creating a sort of “ball” around a sequence in protein space, then profiles increase the volume occupied by both training and test sequences, increasing the chance of an overlap, i.e. of information bleeding between training and test sets (Figure 1), thereby reducing the quality and fairness of the evaluation. Thus here we set out to study these subtle effects and consider the possibility that the observed gain in accuracy could be at least in part due to an evaluation bias – the bias that results from having sequence-based redundancy reduction protocols for evaluating profile-based predictors.

**Figure 1:**
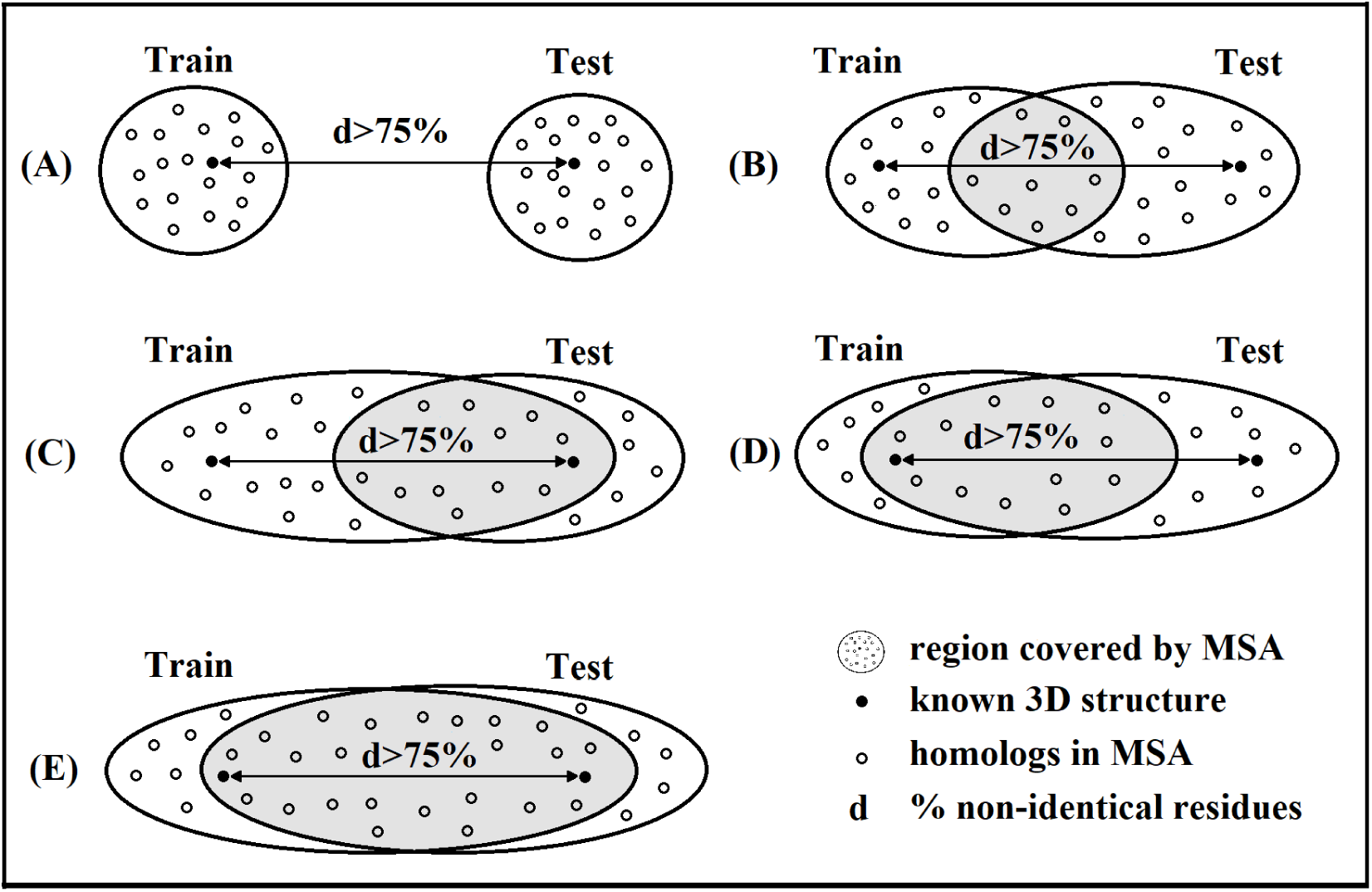
Illustration of various cases of protein space coverage obtained when using evolutionary profiles, derived from multiple sequence alignments (MSA), to train and evaluate a predictor on two proteins sharing less than 25% sequence identity. (A) corresponds to the case initially anticipated in the early 90s (Figure 2 of Rost and Sander, 1993). Other cases observed in this study are depicted by (B), (C), (D), and (E). Profiles lead to different kinds of overlap between training and test sequences.

### 1.1. Three decades of profile-based predictors

The transition from sequence-based to profile-based predictors occurred after a series of landmark studies in the 80s and 90s revealing on one side the relationship between sequence and structure (Doolittle, 1981; Chothia and Lesk, 1986; Sander and Schneider, 1991), and on the other side, providing fast alignment methods to detect putative homologous proteins in large un-annotated protein databases (Sankoff and Kruskal, 1983; Lipman and Pearson, 1985; Altschul et al., 1990, 1997). It became clear at this point that a single protein sequence was sufficient to retrieve information about the entire protein family and its evolution. Evolutionary profiles, calculated from multiple sequence alignments (MSA) of the putative homologous proteins and expressed in the form of amino acid frequencies at each alignment position or position-specific scoring matrices (PSSM), were rapidly selected as a solution to represent and incorporate this newly available information into prediction systems in place of the previously used sequence-based features. The resulting gain in accuracy observed in these studies was striking. For instance, sequence-based secondary structure predictors available in the early 90s with an estimated accuracy between 60% and 65% (Qian and Sejnowski, 1988) were rapidly replaced by a new generation of profile-based predictors with an estimated accuracy between 70% and 76% (Rost and Sander, 1993; Jones, 1999). Since then, predictors have kept improving thanks notably to more sophisticated prediction methods and larger databases (Yang et al., 2016; Jiang et al., 2017) but still belong to the same generation of profile-based predictors. The ∼10% gain in accuracy initially observed is still visible nowadays as recently showed in Heffernan et al., 2018 and Torrisi et al., 2019. Similar trends can be observed for predictors beyond secondary structure to the point that today most state-of-the-art predictors include evolutionary profiles in their input representations.

### 1.2 Assessing predictors: redundancy versus evolutionary profiles

The general evaluation protocol used in the field to assess the accuracy of profile-based predictors was proposed in Rost and Sander, 1993. With this protocol, training and test proteins are selected from the Protein Data Bank (PDB, Gilliland et al., 2000), or from databases directly derived from the PDB like SCOPe (Chandonia et al., 2013) or CATH (Orengo et al., 2016), via a redundancy reduction step performed at the sequence identity level using tools like PSI-BLAST (Altschul et al., 1997), CD-HIT (Godzik and Li, 2006), or Pisces (Wang and Dunbrack, 2003). This protocol was suggested based on the assumption that a protein sequence sharing less than 25% identity with the protein sequences used to train the predictor is a suitable independent test protein to evaluate a profile-based predictor, an assumption clearly visible from Figure 2 of Rost and Sander, 1993 (depicted by case (A) in Figure 1 of this study). However, we have known for a long time that it isn’t particularly unlikely that such a protein may belong to the same family as some of the training proteins (Sander and Schneider, 1991; Brenner et al., 1998; Rost, 1999; Sauder et al., 2000). Such cases would inevitably result in high levels of similarity between the corresponding profiles despite the low level of sequence identity (cases (B) to (E) in Figure 1), and may be the reason for much of the gain in accuracy of profile-based predictors. Likewise, overall differences in accuracy between predictors could result from there being different proportions of cases (A) to (E) in Figure 1 between their training sets and the sets they are tested on.

**Figure 2:**
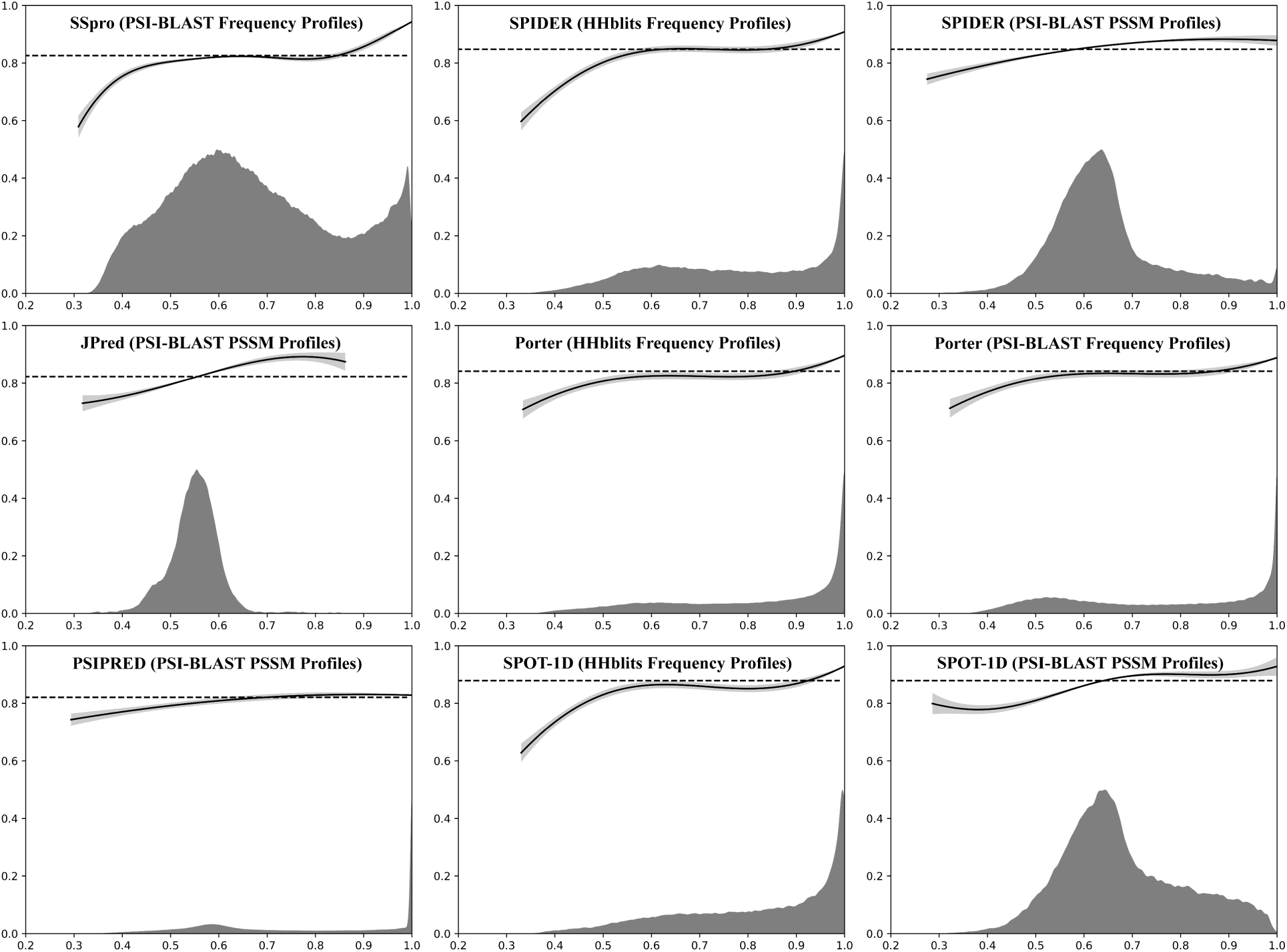
Evaluation results for the six profile-based secondary structure predictors considered in this study on their own test datasets (Porter’s test set for SSpro and PSIPRED) following the protocol described in Section 2.5. The legend for each plot is as follows: (a) the x-axis represents the profile similarity level calculated as indicated in Section 2.1, (b) the y-axis represents the estimated predictor accuracy, (c) the GPR-learned functions interpolating the numerous observations at each profile similarity level are drawn using continuous black curves with 95% confidence intervals drawn around the curve in grey color, (d) the predictor’s average accuracy on the entire test dataset is depicted using discontinuous horizontal black lines, and (e) the frequency of profile fragments observed at each profile similarity level is depicted by plain grey areas in the lower parts of the plots, re-scaled for improved visibility.

### 1.3 A profile-induced evaluation bias?

To better understand the mechanism by which evolutionary profiles contribute to a predictor’s accuracy, we first designed and implemented a simple evaluation protocol to assess the accuracy of a predictor as a function of a profile similarity measure between training and test proteins, calculated using a sliding windows of fixed length. We then applied this protocol to six state-of-the-art profile-based secondary structure predictors using both their respective training and test datasets, whenever available, and a separate test dataset specifically prepared for these predictors. The results of this first set of experiments confirm our initial suspicions, notably: (1) high levels of profile similarity between training and test examples can be observed despite the low level of sequence identity between the corresponding proteins; (2) the accuracy of the predictors is strongly correlated to the level of profile similarity; and (3) high levels of profile similarity are necessary for the predictors to perform significantly better than sequence-based predictors. We then confirm that the redundancy between training and test datasets introduced by the use of evolutionary profiles is the consequence of a larger than initially anticipated coverage of the protein space by these profiles (illustrated in Figure 1). We finally show these conclusions are not specific to secondary structure predictors by showing that comparable results are obtained when using other kinds of predictors. Take together, these results suggest that the use of evolutionary profiles introduces evaluation biases within the current protocols, and that the level and distribution of profile similarity between training and test sets should be explicitly considered when evaluating and comparing different predictors.

## 2. Methods

### 2.1. Profile similarity measure

Measuring the similarity between evolutionary profiles is a problem usually addressed in the context of identifying homologous regions of proteins based on their respective evolutionary profiles. For instance, HHblits (Remmert et al., 2011) detects homologous sequences by performing numerous pairwise alignments of profile HMMs. While remarkably efficient at identifying homologous protein regions, these approaches are more complex and costly to implement than needed in this study. Indeed, the similarity we wanted here is a numerical similarity between the training and test examples of a predictor, independent from the biology, fast to compute on any pair of profiles, and with no need to consider possible insertions in the profiles. We also wanted this measure to be calculated between short protein regions of fixed length in order to address the likely variations of profile similarity along the protein sequences and to limit this measure to a very rough approximation of the information used by the predictors to make a prediction for a given sequence position. We selected a window length of 30 amino acids in our experiments as a compromise between various considerations such as the actual window sizes used by the predictors and the significance of the measure for the selected length. This simplified the problem to measuring the similarity between two 30×20 numerical matrices that we further simplified by flattening the matrices into vectors of dimension 600 without any loss of generality since the matrix structure is not exploited by any of the predictors considered in this study. We selected the cosine similarity measure between the vectors of dimension 600 in our experiments for its desirable properties including notably its low complexity, amplitude independence, and [0, 1] boundaries for positive spaces ([−1, 1] otherwise). For any profile window A of length 30 in a test protein, we therefore define the similarity of A with the training dataset as the maximum cosine similarity value calculated between A and all profile windows of length 30 in the corresponding training dataset.

### 2.2. Profile-based secondary structure predictors and corresponding training/test datasets

We selected six widely used protein secondary structure predictors such that (1) evolutionary profiles constitute the largest part of their input features and (2) the training and test datasets used in the corresponding studies were prepared using a 25% sequence identity threshold. Namely, we used:

- SSpro (Pollastri et al., 2002; Cheng et al., 2005; Magnan and Baldi, 2014)
- JPred (Cuff and Barton, 2000; Drozdetskiy et al., 2015)
- PSIPRED (Jones, 1999; Buchan and Jones, 2019)
- SPIDER (Heffernan et al., 2015, 2017)
- Porter (Pollastri and McLysaght, 2004; Torrisi et al., 2019)
- SPOT-1D (Hanson et al., 2018)

For each predictor, we also collected the corresponding training and test datasets whenever made available by the authors and retrieved the protein databases used by these predictors to generate the profiles. A summary of the software releases, datasets, and profile generation methods used in this study is provided in Table 1. Note however that:

**Table 1:** Description of the profile-based predictors used in this study. The reported release date for each predictor corresponds to the date the models were trained whenever available, to the release date of the software otherwise. The reported date for each dataset is such that all proteins in the corresponding training set had their structure available prior to that date. The protein database release reported in the last column is the release currently used by the online version of the predictors whenever the information was available, a close match with the predictor’s release date otherwise.

| Predictors |  | Datasets |  |  | Profiles |  |  |  |
| --- | --- | --- | --- | --- | --- | --- | --- | --- |
| Name & Release | Date | Date | Training | Test | Method | Type | Database | Release |
| SSpro, ACCpro 5.1 | 10/2013 | 08/2013 | 5,772 | - | PSI-Blast | Frequency | UniRef50 | 06/2015 |
| JPred 4 | 12/2014 | 07/2014 | 1,348 | 149 | PSI-Blast | PSSM | UniRef90 | 07/2014 |
| PSIPRED 4.02 | 03/2016 | 03/2016 | 10,739 | - | PSI-Blast | PSSM | UniRef90 | 05/2016 |
| SPIDER 3 | 10/2016 | 06/2014 | 4,590 | 1,199 | PSI-Blast | PSSM | UniRef90 | 05/2016 |
|  |  |  |  |  | HHblits | Frequency | UniProt20 | 02/2016 |
| Porter, PaleAle 5 | 03/2018 | 12/2014 | 15,753 | 3,154 | PSI-Blast | Frequency | UniRef90 | 05/2016 |
|  |  |  |  |  | HHblits | Frequency | UniProt20 | 02/2016 |
| SPOT-1D | 08/2018 | 02/2017 | 10,029 | 1,213 | PSI-Blast | PSSM | UniRef90 | 04/2018 |
|  |  |  |  |  | HHblits | Frequency | UniClust30 | 10/2017 |

- SSpro and Porter were evaluated without using their template-based prediction modules to remove the evaluation bias they would introduce.
- The datasets used for the latest release of PSIPRED were not retained by the authors so we extracted a fairly large representative training set from the PDB snapshot matching with the release date of the predictor using Pisces (Wang and Dunbrack, 2003) with default settings.
- SSpro was originally evaluated using a cross-validation procedure on a large protein dataset. We therefore considered this dataset as being only the training set of the predictor in this study.
- Porter’s test set is suitable for an independent evaluation of SSpro as all the proteins in this dataset were released after June 2017 and share less than 25% sequence identity with any protein in the training set of SSpro. We also used it to test PSIPRED in our experiments despite high redundancy levels with the PSIPRED training set described above.

### 2.3. Sequence-based secondary structure predictor

In order to get a baseline accuracy for sequence-based prediction of the secondary structure in our experiments and compare the performances between sequence-based and profile-based prediction, we used the most recent predictor we found in that category: SPIDER3 single (Heffernan et al., 2018). Other sequence-based predictors were considered during our study but were all systematically outperformed by SPIDER3 single so only comparisons with this predictor’s accuracy are reported here.

### 2.4. Independent test protein dataset

We extracted from the PDB an independent test set of proteins for the seven predictors listed in Sections 2.2 and 2.3 so that all predictors could be evaluated on the same set of proteins with less than 25% sequence identity with any of the proteins used to train all the predictors, i.e. following the evaluation protocol currently used in the field. PDB entries deposited after February 2017, with less than 25% sequence identity with any of the proteins in the seven training datasets (sequence identity was estimated using PSI-BLAST), and not violating any of the predictor-specific restrictions on the protein sequences (minimal and maximal sequence lengths, no non-standard or unknown amino acids) were selected. The process resulted in 409 such proteins. We name the corresponding protein dataset PDB409.

### 2.5. Profile similarity based evaluation protocol

We implemented the protocol described below to assess the accuracy of a profile-based predictor as a function of the similarity level calculated between the profiles of its training and test proteins.

1. Evolutionary profiles are extracted for each training and test protein following the methods reported in Table 1, i.e. using the same tools and protein databases as the ones originally used to train each predictor.
2. Predicted secondary structures are obtained for each test protein by running the corresponding predictor with the same protein database as the one used to generate the profiles in the previous step.
3. Prediction accuracy and profile similarity level with the proteins in the training dataset are calculated for each possible fragment of length 30 in the test proteins using a sliding window approach. Profile similarity levels are calculated as described in Section 2.1. For the three predictors using two different profiles in input, we consider each profile separately and report the results for both types of profiles.
4. We use the implementation of the Gaussian Process Regression (GPR) method publicly available in the scikit-learn python library to interpolate the numerous pairs (profile similarity level, accuracy) obtained during the previous step. The resulting functions are used to estimate and plot the predictors accuracy at each profile similarity level.

## 3. Results

The results obtained following this protocol are reported individually for each of the six profile-based secondary structure predictors in Figure 2 when evaluating the predictors using their own test sets and in Figure 3 when evaluating them using the PDB409 dataset. A combined view for each set of results and a comparison with the accuracy of the top-performing sequence-based predictor (SPIDER3 single) on the same test datasets are provided respectively in Figures 4 and 5.

**Figure 3:**
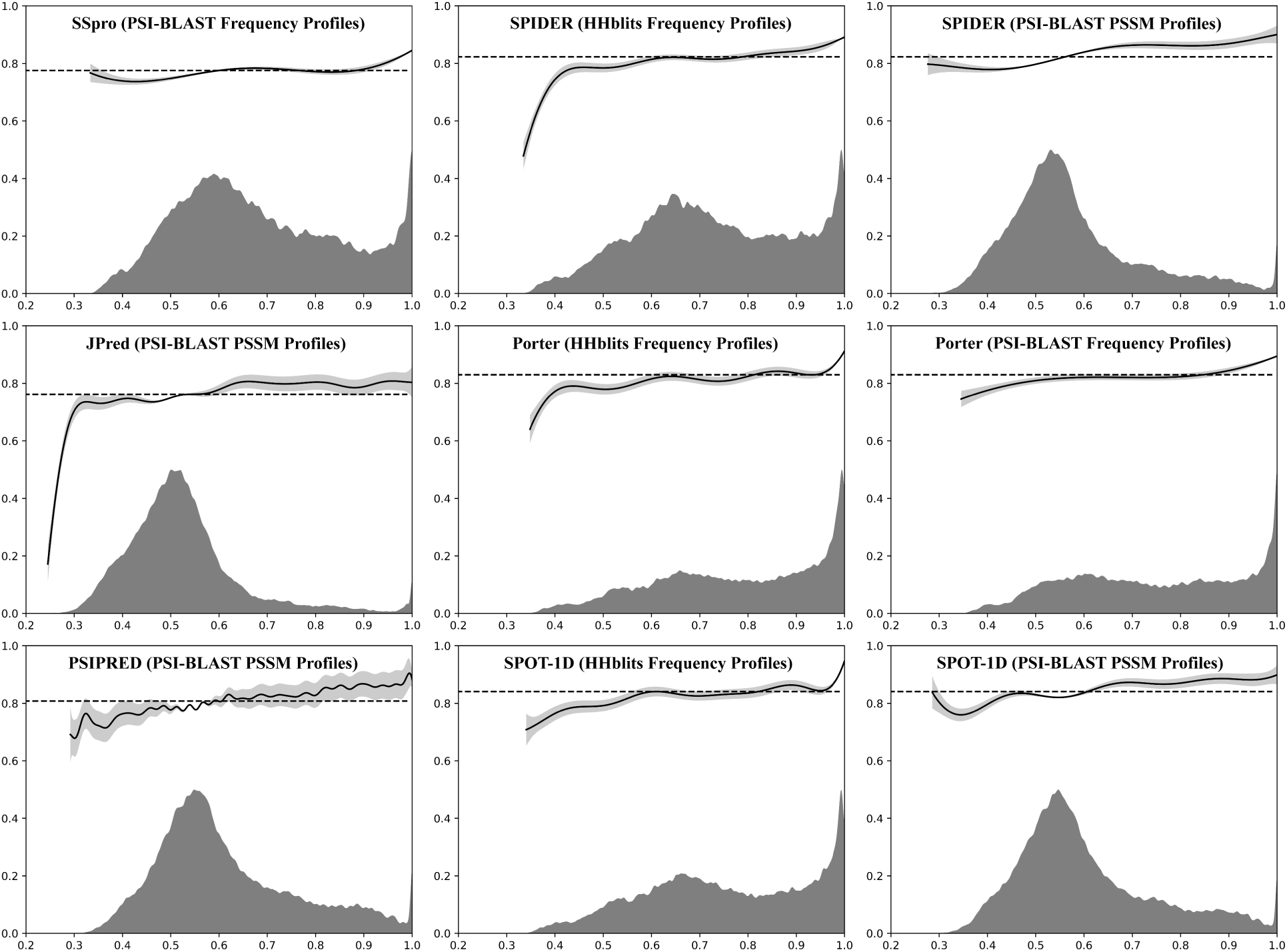
Evaluation results for the six profile-based secondary structure predictors considered in this study on the PDB409 test dataset using the same representation than in Figure 2.

**Figure 4:**
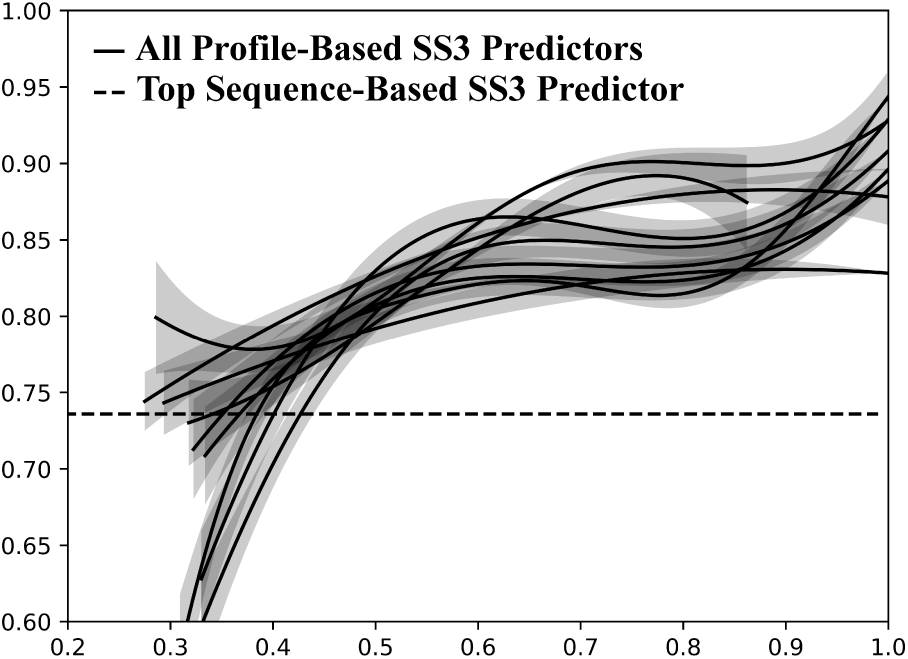
Combined view of the evaluation results provided separately for each profile-based secondary structure predictor in Figure 2, obtained on their own test datasets. Predictor names are omitted for visibility purposes and the average accuracy of SPIDER3 single on the same test sets is indicated for comparison purposes by the discontinuous black line.

**Figure 5:**
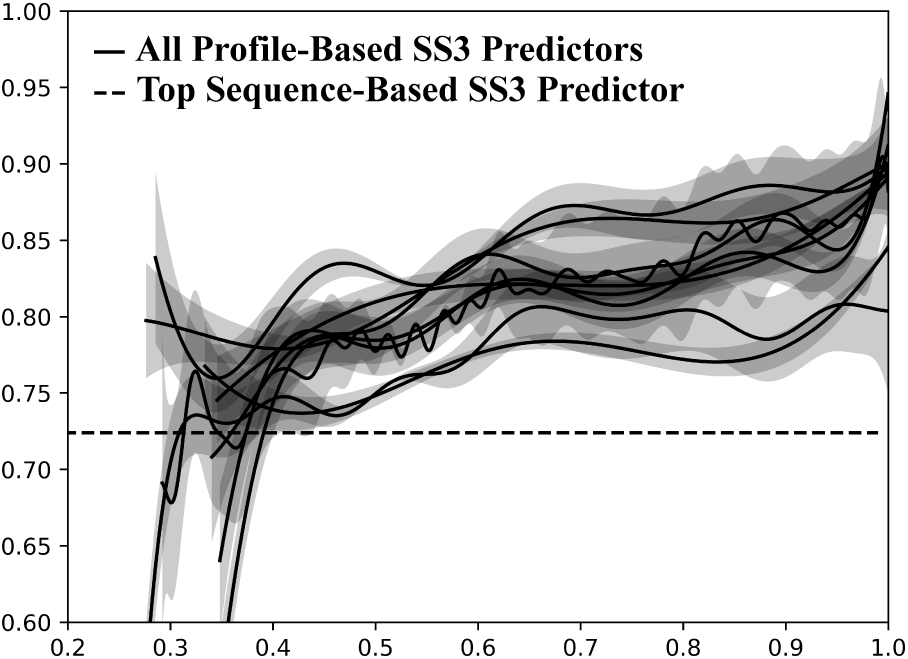
Combined view of the evaluation results provided separately for each profile-based secondary structure predictor in Figure 3, obtained on the PDB409 test dataset. Predictor names are omitted for visibility purposes and the average accuracy of SPIDER3 single on PDB409 is indicated for comparison purposes by the discontinuous black line.

### 3.1. Origin of the redundancy observed when using profiles

The results obtained following the protocol described in Section 2.5 and reported in Figures 2 and 3 show clearly that high frequencies of test profile windows very similar or identical to the training profile windows can be observed in several cases despite the very low level of sequence identity between the proteins. A quick observation of the corresponding MSAs revealed large intersections between the members of each MSA, providing a natural explanation for the high profile similarity values obtained on these examples. The various cases we observed (illustrated in Figure 1) suggest that the protein space covered by the members of an MSA may be much larger than initially anticipated in the 90s (depicted by case (A) in Figure 1) and could be a major factor influencing the overall accuracy of the profile-based predictors developed during the last decades.

We test this assumption by evaluating both a profile-based predictor on sequences and a sequence-based predictor on profiles. On one side, the experiment aims to check if the high accuracy of a profile-based predictor is strongly dependent on the large intersections mentioned above by observing the change in accuracy resulting from removing these intersections. On the other side, the experiment aims to check if the low accuracy of sequence-based predictors is improved by adding such intersections. We trained the two predictors using the same encoding for both sequences and profiles so that the two types of input data could be used to evaluate the predictors. Sequences were represented as frequency profiles containing only 0 and 1 values as commonly done by sequence-based predictors. No extra features were used in these experiments. The dataset and methods used to train the two predictors are not detailed here but are identical for each predictor and closely follow the protocols used in Torrisi et al., 2019. The PDB409 dataset described in Section 2.4 is used to evaluate the trained models both on sequences and profiles generated using HHblits. The accuracy of each model on both types of input data is reported in Table 2.

**Table 2:** Observed accuracy of the profile-based and sequence-based predictors trained as described in Section 3.1 on the PDB409 dataset.

|  | **Tested on sequences** |
| --- | --- |
| **Trained from sequences** | 72.3% |
| **Trained from profiles** | 68.6% |
|  | **Tested on profiles** |
| **Trained from sequences** | 74.5% |
| **Trained from profiles** | 81.5% |

### 3.2. Profile-based prediction of other structural features

As mentioned in introduction, the use of evolutionary profiles to predict structural features of a protein is not limited to the secondary structure prediction problem. We performed the same analysis on different kinds of predictors to make sure that the main results of this study are not specific to the profile-based prediction of the secondary structure. All these experiments led to highly similar results and conclusions so we decided to report only some of these results for the three prediction problems listed below.

- Secondary Structure (8-class)
- Relative Solvent Accessibility (2-class)
- Torsion Angles (14-class)

We selected three predictors for each prediction problem among the ones already used during the previous experiments as most of these predictors are also trained to predict other structural features than the secondary structure 3-class, occasionally distributed under a different name. Datasets and profile generation methods reported in Table 1 are also valid for the corresponding predictors evaluated in this experiment. Evaluation results are reported in Figure 6 and were obtained by evaluating each predictor on its own test dataset similarly to the results reported in Figure 2.

**Figure 6:**
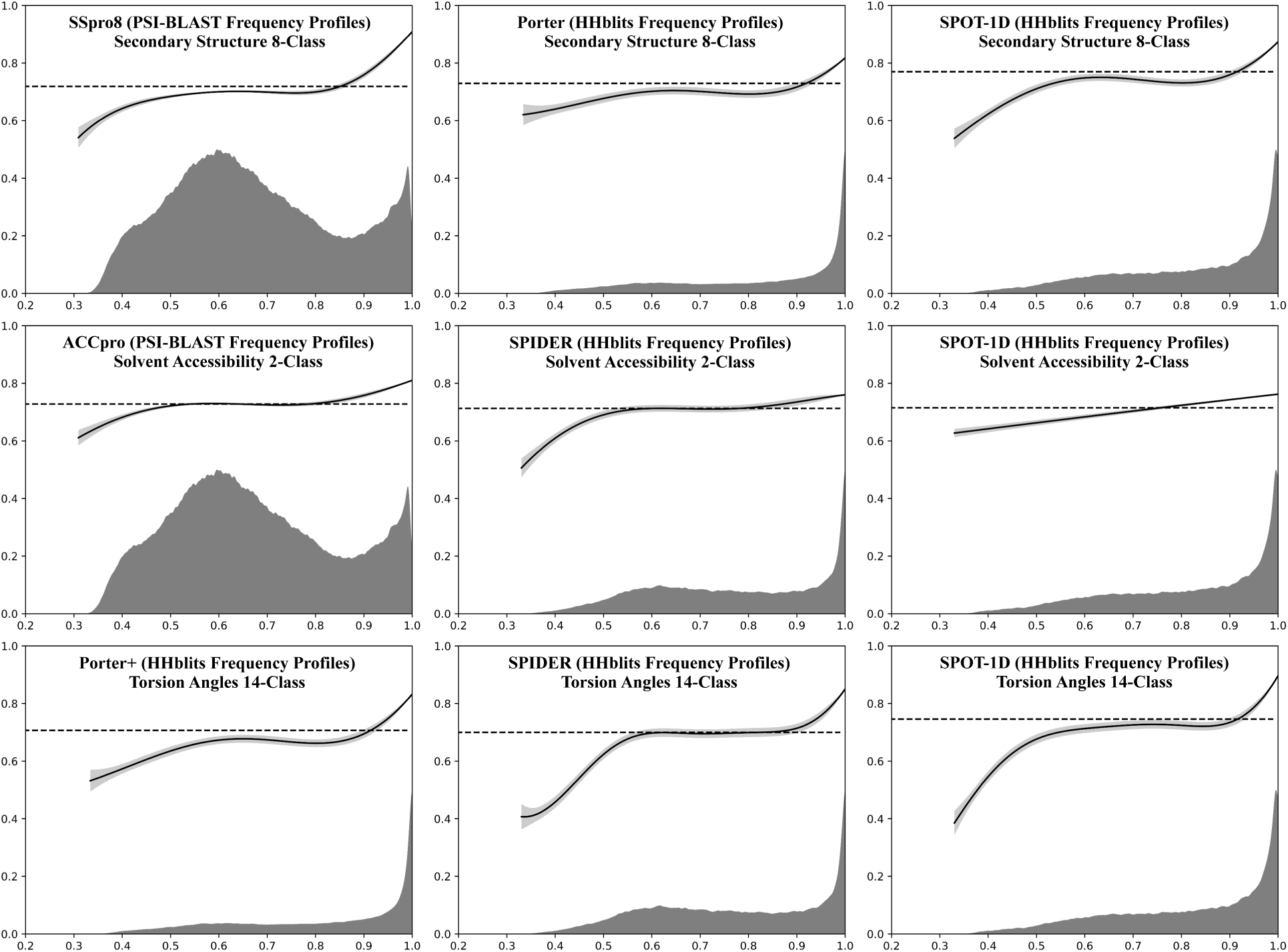
Evaluation results for profile-based predictors of structural features other than the secondary structure 3-class on their own test dataset using the same representation than in Figure 2.

## 4. Discussion

Studying the accuracy of profile-based predictors as a function of the profile similarity between training and test datasets provides interesting results at two different levels. On one side, it reveals the mechanism by which evolutionary profiles increases the prediction accuracy. On the other side, it reveals their use introduce a fairly significant evaluation bias with the protocols used in the field. Both aspects are discussed separately below.

### 4.1. How profiles improve prediction accuracy

The common belief that evolutionary profiles are more informative, compared to single sequences, in structural feature prediction must be qualified in light of the results obtained in this study. All the predictors evaluated in our experiments show a clear correlation between their prediction accuracy and the level of profile similarity. It is striking to see how similarly all these predictors perform when looking at the results reported in Figures 4 and 5. All predictors perform overall poorly on low profile similarity fragments with, in some cases, a level of accuracy which is even below the level achieved by pure sequence-based predictors. And all predictors improve steadily as the profile similarity level increases. A proper evaluation of a predictor should use test proteins that are unrelated to the training proteins. We have seen that this is not systematically the case with current protocols where variable levels of redundancy are observed as a result of profiles being calculated from alignments of homologous sequences that contain identical subsets. The results reported in Table 2 show that profile-based predictors are unable to sustain their performances without this redundancy. This result is also visible when performing the opposite experiment, i.e. adding some redundancy between the training and test sequences of a sequence-based predictor, by replacing test sequences with profiles, which ends up boosting the predictor’s accuracy. Taken together, these results show that the ∼ 10% accuracy difference between profile-based and sequence-based predictors is in part due to large quantities of highly similar or identical profiles in the training and test datasets, a regime where machine learning methods are naturally expected to be more accurate.

### 4.2. Consequences for current evaluation protocols

The results obtained in this study are also evidence that the current evaluation protocols used by the community are not adequate to: (1) reliably assess the accuracy of a profile-based predictor; and (2) compare profile-based predictors. Indeed, results reported in Figures 2, 3, and 6 show that the mean accuracy of a profile-based predictor will strongly depend on the abundance of profile fragments highly similar between training and test sets. From the same results, one can also see that this abundance is not constant at all when using a 25% sequence identity threshold to reduce the redundancy between training and test datasets, leading to important variations of the estimated accuracy of a predictor from a test set to another (up to 6.1% between the results reported in Figure 2 and 3 for instance). Even comparing the predictors on identical datasets would not solve this issue as the abundance of high similarity profile fragments is also dependent on other factors such as: the method used to generate the profiles; the protein database used to find homologous proteins; and even the type of profiles used in input of the predictors. An evaluation protocol assessing the accuracy of a profile-based predictor as a function of the similarity level between training and test profiles, such as the one implemented in this study, can actually solve this issue since predictors can then be compared at each profile similarity level. It also presents several major advantages: (1) it removes the need to compare predictors on the same datasets; (2) it naturally separates easy test examples from hard ones, making unnecessary to even decide if a protein is a suitable test example or not; and (3) it is simple to implement and does not require a lot of computation time. If a protocol, such as the one described here, is used to assess and compare predictors, the need to separate training and test sets using a strict, but arbitrary, sequence identity threshold becomes redundant, leading to the possibility of adopting larger training sets and designing predictors that have a higher accuracy over a larger portion of the protein space.

### 4.3. Software availability

An implementation of the evaluation protocol proposed in this study, named EVALpro, is available for download, with a full documentation, from the SCRATCH suite (Cheng et al., 2005) at: http://scratch.proteomics.ics.uci.edu. It can be used to either reproduce the analysis presented here or, more generally, to evaluate other profile-based predictors and training/test sets.

## 5. Conclusion

By probing the mechanisms behind the increase in average accuracy of profile-based predictors versus sequence-based predictor, we have shown that this increase largely relies on the presence and level of redundancy introduced by the use of evolutionary profiles. This redundancy produces evaluation biases when current evaluation protocols are used. Despite these results, and somewhat paradoxically, the usefulness of including evolutionary profiles in the predictors’ inputs remains unchanged. This is for the same reasons, and with the same limitations, as using template-based prediction methods when templates are available in structural databases. In both cases, a significant improvement in accuracy can be expected; but this improvement occurs only when certain conditions of overlap are met. The continued growth of protein databases benefits profile-based predictors by increasing the number of situations where these favorable overlap conditions occur. Nevertheless, at the opposite end, our work clearly shows that the evaluation protocols used in the field need to be revised to account for the biases associated with these overlaps. In particular, we have shown that measuring average accuracy alone on a protein data set is not particularly meaningful or reliable. Instead, we should measure accuracy as a function of profile similarity. Such a protocol provides a mean for evaluating profile-based predictors, and compare them with each other and with sequence-based predictors, in a fairer way.

